# Cryo-EM of the yeast V_O_ complex reveals distinct binding sites for macrolide V-ATPase inhibitors

**DOI:** 10.1101/2021.11.15.468710

**Authors:** Kristine A. Keon, Samir Benlekbir, Susanne H. Kirsch, Rolf Müller, John L. Rubinstein

**Affiliations:** Molecular Medicine Program, The Hospital for Sick Children, Toronto, Canada M5G0A4; Department of Medical Biophysics, The University of Toronto, Toronto, Canada M5G1L7; Helmholtz Institute for Pharmaceutical Research Saarland (HIPS), Helmholtz Centre for Infection Research (HZI) and Department of Pharmacy, Saarland University Campus, 66123 Saarbrücken, Germany; Department of Biochemistry, The University of Toronto, Toronto, Canada M5S1A8

## Abstract

Vacuolar-type adenosine triphosphatases (V-ATPases) are proton pumps found in almost all eukaryotic cells. These enzymes consist of a soluble catalytic V_1_ region that hydrolyzes ATP and a membrane-embedded V_O_ region responsible for proton translocation. V-ATPase activity leads to acidification of endosomes, phagosomes, lysosomes, secretory vesicles, and the trans-Golgi network, with extracellular acidification occurring in some specialized cells. Small molecule inhibitors of V-ATPase have played a crucial role in elucidating numerous aspects of cell biology by blocking acidification of intracellular compartments, while therapeutic use of V-ATPase inhibitors has been proposed for treatment of cancer, osteoporosis, and some infections. Here, we determine structures of the isolated V_O_ complex from *Saccharomyces cerevisiae* bound to two well-known macrolide inhibitors: bafilomycin A1 and archazolid A. The structures reveal different binding sites for the inhibitors on the surface of the proton-carrying c ring, with only a small amount of overlap between the two sites. Binding of both inhibitors is mediated primarily through van der Waals interactions in shallow pockets and suggests that the inhibitors block rotation of the ring. Together, these structures indicate the existence of a large chemical space available for V-ATPase inhibitors that block acidification by binding the c ring.

## Introduction

Vacuolar-type adenosine triphosphatases (V-ATPases) are large membrane-embedded protein complexes that hydrolyze ATP to pump protons across the membrane. V-ATPases, which are ubiquitous in eukaryotic cells, acidify secretory vesicles, lysosomes, endosomes, and the trans Golgi network (Hinton et al., 2009). In mammals, V-ATPases also secrete protons from osteoclasts (Blair et al., 1989; Qin et al., 2012), kidney intercalated cells (Brown et al., 2009), and some tumour cells (Stransky et al., 2016). The mammalian enzyme includes 16 different subunits, with several subunit possessing isoforms encoded by different genes and expressed in a cell- and tissue-dependent manner (Toei et al., 2010). Mutations in components mammalian V-ATPases may be either embryonic lethal (Sun-Wada et al., 2000) or result in numerous different diseases (Hinton et al., 2009).

V-ATPases are comprised of a catalytic V_1_ region, which is found in the cytosol and hydrolyzes ATP, and membrane-embedded V_O_ region, which is responsible for proton translocation (Vasanthakumar and Rubinstein, 2020). ATP hydrolysis in the V_1_ region induces rotation of a central rotor subcomplex that drives proton translocation through V_O_. Protons are conducted through the V_O_ region by binding to conserved Glu residues on each c, c′ and c′′ subunit, which rotate against subunit a to collect protons from the cytoplasm and deposit them on the other side of the membrane (Mazhab-Jafari et al., 2016). The V_1_ and V_O_ regions can undergo reversible dissociation to regulate V-ATPase activity (Kane, 1995; Sumner et al., 1995), with ATP hydrolysis inhibited in the dissociated V_1_ complex and the dissociated V_O_ complex becoming impermeable to protons (Parra et al., 2000; Zhang et al., 1992). The V_1_ complex contains subunits A_3_B_3_CDE_3_FG_3_H, while the V_O_ complex includes subunits a, d, e, and f, as well as the c ring. In yeast, the c ring has a stoichiometry of c_8_c′c′′ with the subunit Voa1p found within the lumen of the ring (Mazhab-Jafari et al., 2016; Roh et al., 2018). Each c, c′, and c′′ subunit has four transmembrane α helices that form an outer ring of 20 α helices and an inner ring of 20 α helices. Cryo-EM of intact V-ATPases identified three rotational states, known as ‘State 1’, ‘State 2’, and ‘State 3’ (Abbas et al., 2020; Zhao et al., 2015). In each rotational state, the three pairs of A and B subunits in the V_1_ region adopt different conformations, with the rotor subcomplex stopped in a different position. Following separation from V_1_, the V_O_ complex is found in a conformation resembling its conformation in State 3 of the intact V-ATPase (Mazhab-Jafari et al., 2016).

V-ATPase inhibitors have served as essential tools for cell biology research, allowing investigators to block acidification of cellular compartments such as lysosomes and synaptic vesicles (Maxson and Grinstein, 2014). Pharmaceutical inhibition of V-ATPases has been proposed for treatment of numerous diseases (Hayek et al., 2014; Kartner and Manolson, 2014; Moreno et al., 1998; Stasic et al., 2019; Stransky et al., 2016). V-ATPase inhibitors for cancer treatment have shown promise by inhibiting metastasis (Schempp et al., 2014; Wiedmann et al., 2012), decreasing protease activation (Kubisch et al., 2014), and upregulating protective cellular responses (Graham et al., 2013). Due to the role of V-ATPase in the breakdown of bone by osteoclasts (Blair et al., 1989; Hollberg et al., 2002; Teitelbaum, 2000; Tezuka et al., 1994), inhibiting V-ATPases in osteoclasts has been explored for the treatment of osteoporosis (Kartner and Manolson, 2014; Sundquist et al., 1990). As V-ATPase is an essential enzyme in parasites such as *Toxoplasma gondii* (Moreno et al., 1998; Stasic et al., 2019) and fungal pathogens such as *Candida albicans* (Hayek et al., 2014), targeting V-ATPase activity has been suggested for treating infection by these organisms. Further, inhibition of human V-ATPase has been proposed to block infection by viruses that depend on endosomal acidification for entry into cells (Chen et al., 2013; Hu et al., 2018; Sabino et al., 2019; Yeganeh et al., 2015).

The most potent and specific V-ATPase inhibitors include the macrolides bafilomycin A1 (Hanada et al., 1990) (Fig. 1A, *left*), concanamycin A (Páli et al., 2004; Whyteside et al., 2005; Woo et al., 1992) (Fig. 1B, *left*), and archazolid A (Sasse et al., 2003) (Fig. 1C, *left*). Bafilomycins and concanamycins were isolated from *Streptomyces* bacteria, and have 16- and 18-membered macrolide rings, respectively (Kinashi et al., 1984; Werner et al., 1984). Bafilomycin A1 is the smallest of the bafilomycin compounds, with other bafilomycins having higher molecular weights due to additional functional groups (Werner et al., 1984).

**Figure 1.**
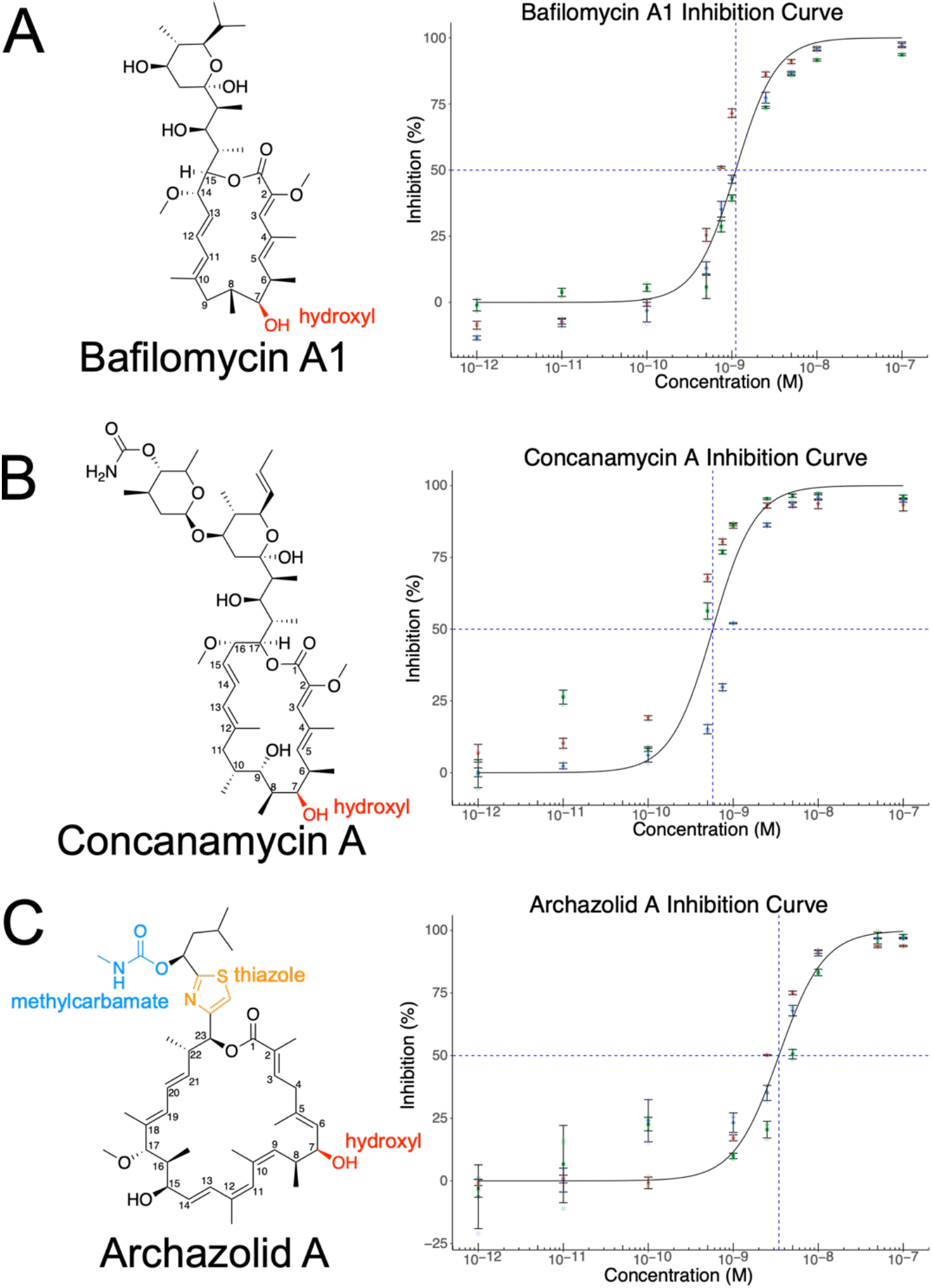
Molecular structure and dose-response curves for V-ATPase inhibitors. **A**, Bafilomycin A structure (*left*) and dose-response curve for ATPases activity of purified yeast V-ATPase (*right*). **B,** Concanamycin A structure (*left*) and dose-response curve with purified yeast V-ATPase (*right*). **C,** Archazolid A structure (*left*) and dose-response curve with purified yeast V-ATPase (*right*).

Concanamycin A is the most potent of the concanamycins (Kinashi et al., 1984). Both macrolide molecules possess a 7′ hydroxyl group on their macrolide ring (Fig. 1A and B, *left*) and have sugar-like moieties branching from the ring. Archazolids were discovered from myxobacteria (Sasse et al., 2003). Members of this class of V-ATPase inhibitors have a 24-membered macrolide ring possessing a 7′ hydroxyl group, a thiazole group, and a methylcarbamate group (Fig. 1C, *left*). Archazolids have been reported to inhibit cancer cell proliferation at concentrations lower than those needed to inhibit V-ATPase activity (Scheeff et al., 2020). V-ATPase inhibition by archazolids decreases levels of the anti-apoptotic protein survivin (Zhang et al., 2015), increases cell stiffness and membrane polarity of cancer cells (Bartel et al., 2017), and differentially regulates AMP-activated protein kinase activity in healthy and cancer cells (Bartel et al., 2019), suggesting potential as a cancer therapeutic.

A 3.6 Å resolution structure of bafilomycin A1 bound to intact bovine V-ATPase revealed the inhibitor on the surface of the c ring between adjacent c subunits (Wang et al., 2021). Inhibitor binding involves extensive contacts between the compound and hydrophobic amino acid side chains from the protein, with Tyr144 (equivalent to Tyr142 in the *S. cerevisae* subunit c) forming a hydrogen bond with the 7′ hydroxyl group of the macrolide ring (Wang et al., 2021). This interaction inhibits V-ATPase activity by blocking the rotation of the c ring at the a-c ring interface. In contrast to these structural studies, the binding site of archazolids was investigated by site-directed mutagenesis, competitive binding with bafilomycin and a fluorescent derivative of N,N-dicyclohexylcarbodiimide (DCCD), which covalently binds the glutamate responsible for proton transport (Bockelmann et al., 2010), and binding studies with archazolid derivatives (Scheeff et al., 2020). Based on this analysis, archazolids were predicted to interact with both the proton-carrying glutamate of the c subunit and the tyrosine residue involved in bafilomycin binding (Bockelmann et al., 2010). However, it is unclear how the VO region accommodates both bafilomycins and archazolids due to the different sizes of the inhibitors. Here we determined structures of the yeast V_O_ complex bound to bafilomycin A1 and to archazolid A. Surprisingly, we find little overlap between the bafilomycin and archazolid binding sites indicating that different inhibitors can bind the V_O_ region in different ways to potently inhibit V-ATPase activity.

## Results and Discussion

We found that intact, detergent-solubilized V-ATPase from *S. cerevisiae* was most active when solubilized, purified, and assayed with the detergent n-Dodecyl-β-D-Maltopyranoside (DDM), with a specific activity of 5.5 ± 0.9 μmol ATP mg^−1^ min^−1^ (± s.d, n=3 independent measurements, each using a different batch of purified protein). In comparison, the specific activity of the enzyme with 0.002% (w/v) glyco-diosgenin (GDN), a detergent that has been particularly useful for structural studies of V-ATPase (Vasanthakumar et al., 2019), had only ~3% of the specific activity of the enzyme in DDM. We next measured dose-response curves for enzyme activity with different concentrations of inhibitors with the DDM-solubilized enzyme in an assay buffer containing DDM. In these assays, 50% inhibition of enzyme activity occurred at inhibitor concentrations comparable to the nanomolar concentration of enzyme needed to perform the assays. Consequently, the measured IC_50_ values (± s.d., n=3 independent titrations for each inhibitor, each titration using a different batch of protein and inhibitor dilution series) provide an upper bound only on the true IC_50_ values for the different inhibitors: 1.11 ± 0.39 nM with a Hill coefficient of 1.82 ± 0.30 for bafilomycin A1 (enzyme concentration measured at 4.7 to 6.4 nM), 0.58 ± 0.41 nM with a Hill coefficient of 1.75 ± 0.45 for concanamycin A (enzyme concentration measured at 1.8 to 4.7 nM), and 3.44 ± 1.99 nM with a Hill coefficient of 1.65 ± 0.28 for archazolid A (enzyme concentration measured at 0.79 to 1.9 nM). The IC_50_ for bafilomycin A1 and concanamycin A were previously reported to have similar low nanomolar IC_50_ values (Wang et al., 2021, 2005) with archazolid A having a somewhat higher IC_50_ than bafilomycin and concanamycin (Bockelmann et al., 2010; Huss et al., 2005). The Hill coefficient being greater than one could indicate that inhibitor binding to the c ring occurs more readily when the c ring’s rotation is blocked by previous binding of an inhibitor molecule to the complex.

To understand how these inhibitors of different sizes result in comparably potent inhibition of the enzyme, we investigated the structure of the *S. cerevisiae* V_O_ complex bound to bafilomycin A1 and archazolid A. Following starvation-induced dissociation of V-ATPase (Mazhab-Jafari et al., 2016), the V_O_ complex was purified and partially exchanged into GDN (see methods), which is amenable to high-resolution structural studies (Guo et al., 2017; Vasanthakumar et al., 2019). Inhibition of intact V-ATPase activity by GDN likely occurs due to binding of the sterol moiety of GDN to the c ring (Vasanthakumar et al., 2019). We reasoned that this low-affinity interaction would not interfere with the high-affinity binding of inhibitors, while the improved resolution from cryo-EM with GDN was critical for interpretation of how inhibitors bind. The V_O_ preparation at ~0.1 to 0.2 mg/mL protein was mixed with a ~150- to 200-fold molar excess of inhibitor before concentrating the sample to 3 to 6 mg/ml and subjecting it structure determination by cryo-EM as described previously (Mazhab-Jafari et al., 2016; Vasanthakumar et al., 2019) (Fig. S1, Table 1). As expected, the resulting V_O_ structures were found in rotational State 3, as seen previously with the V_O_ complex (Mazhab-Jafari et al., 2016; Roh et al., 2020, 2018; Vasanthakumar et al., 2019) (Fig. S1). The overall resolutions of the maps, 3.2 Å for the bafilomycin-bound structure and 2.8 Å for the archazolid-bound structure, allowed for construction of atomic models of the complexes (Fig. S2). While side chains were modelled for both the membrane-embedded and soluble regions of the complex, resolution was better in the membrane-embedded regions (Fig. S1B), consistent with prior V_O_ structures (Mazhab-Jafari et al., 2016; Roh et al., 2020; Vasanthakumar et al., 2019).

**Table 1:** Cryo-EM Data Collection, refinement, and validation statistics.

|  | Vo + Bafilomycin A1 | Vo + Archazolid A |
| --- | --- | --- |
| **Data collection and processing** |  |  |
| Magnification | 75,000 | 75,000 |
| Voltage (kV) | 300 | 300 |
| Electron exposure (e-/Å<sup>2</sup>) | 39 | 45 |
| Defocus range (µm) | 0.65-2.76 | 0.37-2.55 |
| Pixel Size (Å) | 1.03 | 1.03 |
| Symmetry imposed | C1 | C1 |
| Map resolution (Å) | 2.8 | 3.2 |
| FSC threshold | 0.143 | 0.143 |
| Map resolution range (Å) | 2.6-3.8 | 2.4-3.6 |
| **Refinement** |  |  |
| Initial model used (PDB code) | Vo + Archazolid A Model | 6M0R |
| Model Resolution (Å) | 3.3 | 2.9 |
| FSC threshold | 0.5 | 0.5 |
| Map sharpening *B* factor (Å<sup>2</sup>) | -135.7 | -126.9 |
| Model composition |  |  |
| Non-hydrogen atoms | 22,497 | 22,532 |
| Protein residues | 2915 | 2915 |
| Ligands | 9 | 8 |
| *B* factors (Å<sup>2</sup>) |  |  |
| Protein | 50.15 | 40.32 |
| Ligand | 46.56 | 31.19 |
| R.m.s. deviations |  |  |
| Bond lengths (Å) | 0.007 (5) | 0.006 (6) |
| Bond angles (°) | 1.464 (62) | 1.234 (24) |
| Validation |  |  |
| MolProbity score | 1.50 | 1.46 |
| Clashscore | 8.41 | 8.45 |
| Poor rotamers (%) | 1.15 | 0.3 |
| Ramachandran plot |  |  |
| Favoured (%) | 98.20 | 98.61 |
| Allowed (%) | 1.74 | 1.32 |
| Disallowed (%) | 0.07 | 0.07 |

### Bafilomycin binding to V-ATPase occurs mainly due to van der Waals interactions

In the bafilomycin A1-bound map, nine densities corresponding to bafilomycin are resolved (Fig. 2A, *yellow densities*, Fig 2B and C, *yellow models,* Fig. S3). Seven of these densities are wedged between the second and fourth transmembrane α helices (TM2 and TM4) of adjacent c subunits (Fig. 2D, Fig S3A to G). We name these sites the “c-c” sites because they involve two adjacent c subunits. The eighth bafilomycin density is between TM2 of subunit c_(8)_ and TM4 of the adjacent c′ subunit, which we name the “c-c′” site (Fig. 2E, Fig S3H), while ninth is in the “c′-c′′” site between TM2 from the c′ subunit and TM5 of the c′′ subunit (Fig. 2F, Fig S3I). The bafilomycin densities in c-c sites are positioned sufficiently close to the conserved Tyr142 of subunit c to accommodate the hydrogen bond with the 7′ hydroxyl group from bafilomycin that was proposed previously (Wang et al., 2021) (Fig. 2D, *circled in blue*). Similarly, the bafilomycin in the c-c′ site is sufficiently close to Tyr150 of the c′ subunit to accommodate an equivalent hydrogen bond (Fig. 2E, *circled in blue*). Even with the improved resolution of the map compared to the earlier structure, the orientation of the macrolide ring is ambiguous, with a model-to-map correlation of 0.57 and 0.58 for the two possible 180° rotations of the macrolide ring. However, the orientation shown allows formation of the hydrogen bonds with Tyr142 from subunit c or Tyr150 from subunit c′. The ninth bafilomycin molecule, bound at the c′-c′′ site, shows the inhibitor at lower occupancy than the two other types of site (Fig. 2F). With the standard deviation of the map normalized to σ=1, bafilomycins in the c-c sites become visible at σ=6 to 8 and bafilomycin in the c-c′ site becomes visible at σ=6, while bafilomycin in the c′-c′′ site becomes visible at σ=4. Subunit c′′ lacks a Tyr residue at an appropriate position for forming a hydrogen bond with the 7′ hydroxyl group from bafilomycin. In the previous lower-resolution structure (Wang et al., 2021) bafilomycin was described as not being able to bind the c′-c′′ site at all due to the absence of the Tyr subunit. The presence of bafilomycin in all three different types of site, but with the highest occupancy at the c-c and c-c′ sites, suggests that the hydrogen bond stabilizes binding, but van der Waals contacts due to close interactions between the inhibitor and c ring are sufficient for at least low-affinity bafilomycin binding. This hypothesis is consistent with prior studies that demonstrated that the affinity of bafilomycin analogues for V-ATPase is decreased, but not eliminated, upon substitutions of the 7′ hydroxyl group on the macrolide ring (Gagliardi et al., 1999, 1998). Close contact and the resulting van der Waals interactions between bafilomycin and the protein occur due to Phe51, Ile54, Val55, Ile58, Gly61, and Ile65 from the first c subunit, and residues Leu131, Ile134, Phe135, and Val138 from the adjacent c subunit (Fig. 2D). Binding at the first of the seven c-c sites also involves contact between bafilomycin and residues Leu780, Ala783, and Met788 from subunit a (Fig. 2D, *residues indicated on green α helix*). Binding at the c-c′ site involves residues Phe51, Ile54, Val55, Ile58, Gly61, and Ile65 from subunit c_(8)_ and residues Leu139, Ile142, Phe143, Val146, Tyr150 from subunit c′ (Fig. 2E) while binding at the c′-c′′ involves residues Met57, Lys58, Ile61, Val64, Gly67, Ile68, and Ile71 from subunit c′ and residues Val186, Ile189, Phe190, and Ile199 from subunit c′′ (Fig. 2F). As described previously (Wang et al., 2021), binding of bafilomycin around the c ring would prevent rotation of the ring, thereby explaining why the high-affinity binding of the compound inhibits V-ATPase activity.

**Figure 2.**
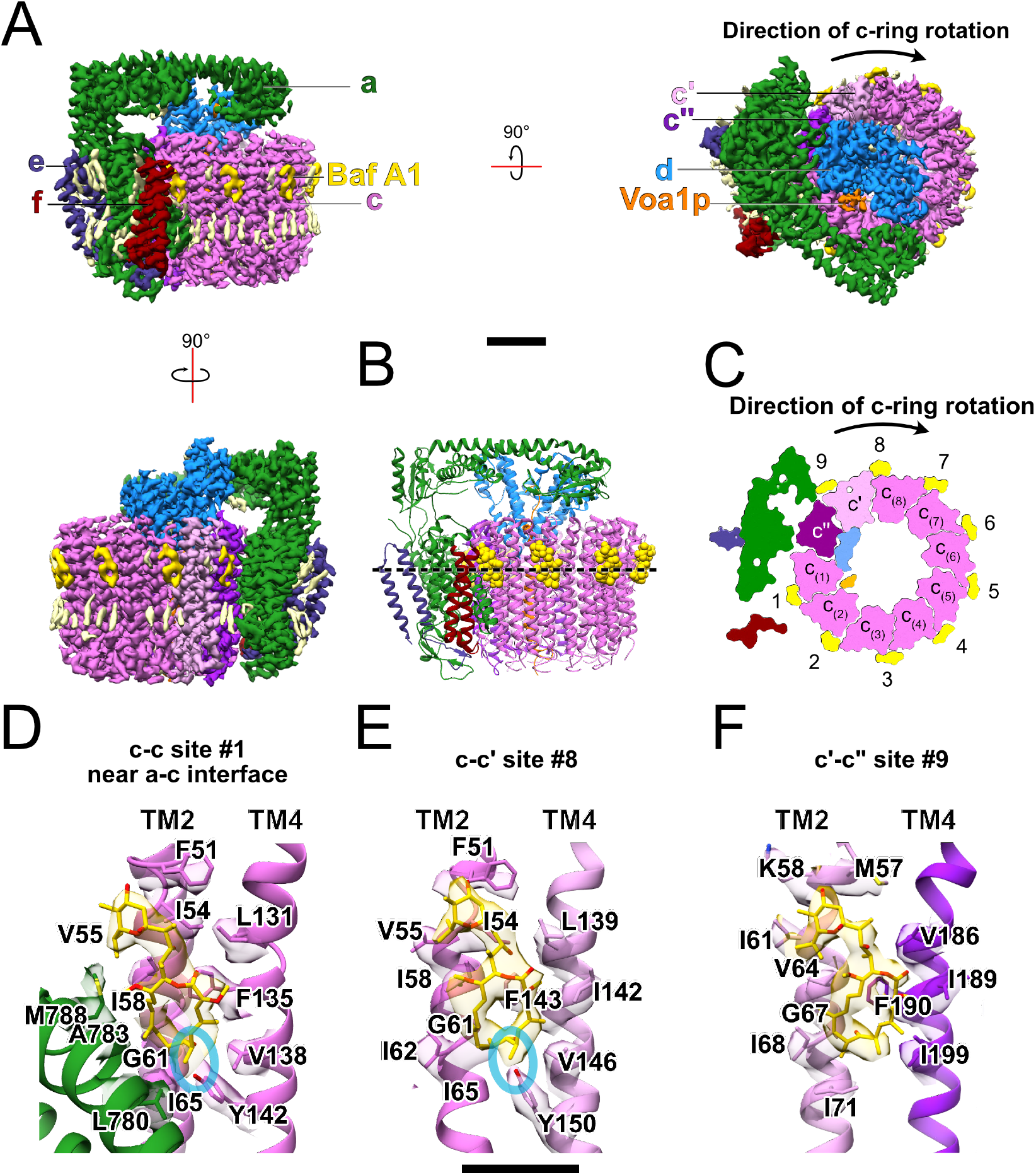
Structure of bafilomycin A1 bound to the yeast the V-ATPase V_O_ region. **A,** Cryo-EM map of bafilomycin A1 bound to the yeast V _O_ region (three views). Scale bar, 25 Å. **B,** Atomic model of bafilomycin A1 bound to the yeast V _O_ complex. **C,** A slice through the atomic model showing locations of nine bafilomycin A1 molecules. **D,** Atomic model showing a c-c site for bafilomycin A1 binding. **E,** Atomic model showing the c-c′ site for bafilomycin A1 binding. **F,**Atomic model showing the c′-c′′site for bafilomycin A1 binding. Scale bar, 10 Å.

### Archazolid binding to V-ATPase involves three α helices on the surface of the c ring

In the archazolid A-bound structure, only eight inhibitor molecules bind the V_O_ complex (Fig. 3A, *light green densities*, Fig. 3B and C, *light green models,* Fig. S4). As with bafilomycin, there are three different types of binding site: six c-c sites (Fig. 3D, Fig. S4A to F), a c-c′ site (Fig. 3E, Fig. S4G), and a c′-c′′ site (Fig. 3F, Fig. S4H). Unlike the bafilomycin binding sites, the archazolid binding sites involve three different transmembrane α helices from the c ring. The c-c sites involves residues Ile58, Met59, Gly61, Ile62, Ile65, Tyr66 on TM2 and Ile134, Glu137, Val138, and Leu141 on TM4 of the first c subunit, and residues Ile134, Phe135, Val138, Leu139, and Tyr142 on TM4 of the second c subunit (Fig. 3D). The c-c′ site involves the same residues as the c-c site from the c_(8)_ subunit, and residues Phe143, Val146, Leu147, and Tyr150 from TM4 of the c′ subunit (Fig. 3E). The c′-c′′ site involves residues Ile64, Met65, Gly67, Ile68, Ile71, Tyr72 on TM2 and Ile142, Glu145, Val146, and Leu149 on TM4 from the c′ subunit, and residues Phe190, Val193, and Leu194 on TM5 of the c′′ subunit (Fig. 3F). Additionally, in the c′-c′′ site, the archazolid molecule interacts with Ile720 of subunit a (Fig. 3F, *residue shown on green α helix*). Inhibitor molecules in the c-c, c-c′, and c′-c′′ sites are positioned to form hydrogen bonds between the 7′ hydroxyl group of archazolid and the proton carrying Glu137 residue of subunit c (or Glu145 of subunit c′ in the c′-c′′ site) and the 15′ hydroxyl group of archazolid and Tyr66 of subunit c (or Tyr72 of subunit c′ in the c′-c′′ site) (Fig. 3D to F, *circled in blue*). In the c-c and c-c′ sites, the carbonyl oxygen from archazolid’s methylcarbamate group and Tyr142 of subunit c or Tyr150 of subunit c′ are positioned to form an additional hydrogen bond (Fig. 3D and E, *circled in red*), for which there is no equivalent in the c′-c′′ site. These interactions between archazolid and the polar and acidic residues were predicted by earlier labelling, site-directed mutagenesis, and pharmacological studies (Bockelmann et al., 2010; Scheeff et al., 2020). Overall, the structure suggests that three hydrogen bonds contribute to archazolid A binding in the c-c and c-c′ sites, and that two hydrogen bonds contribute to archazolid A binding in the c′-c′′ site.

**Figure 3.**
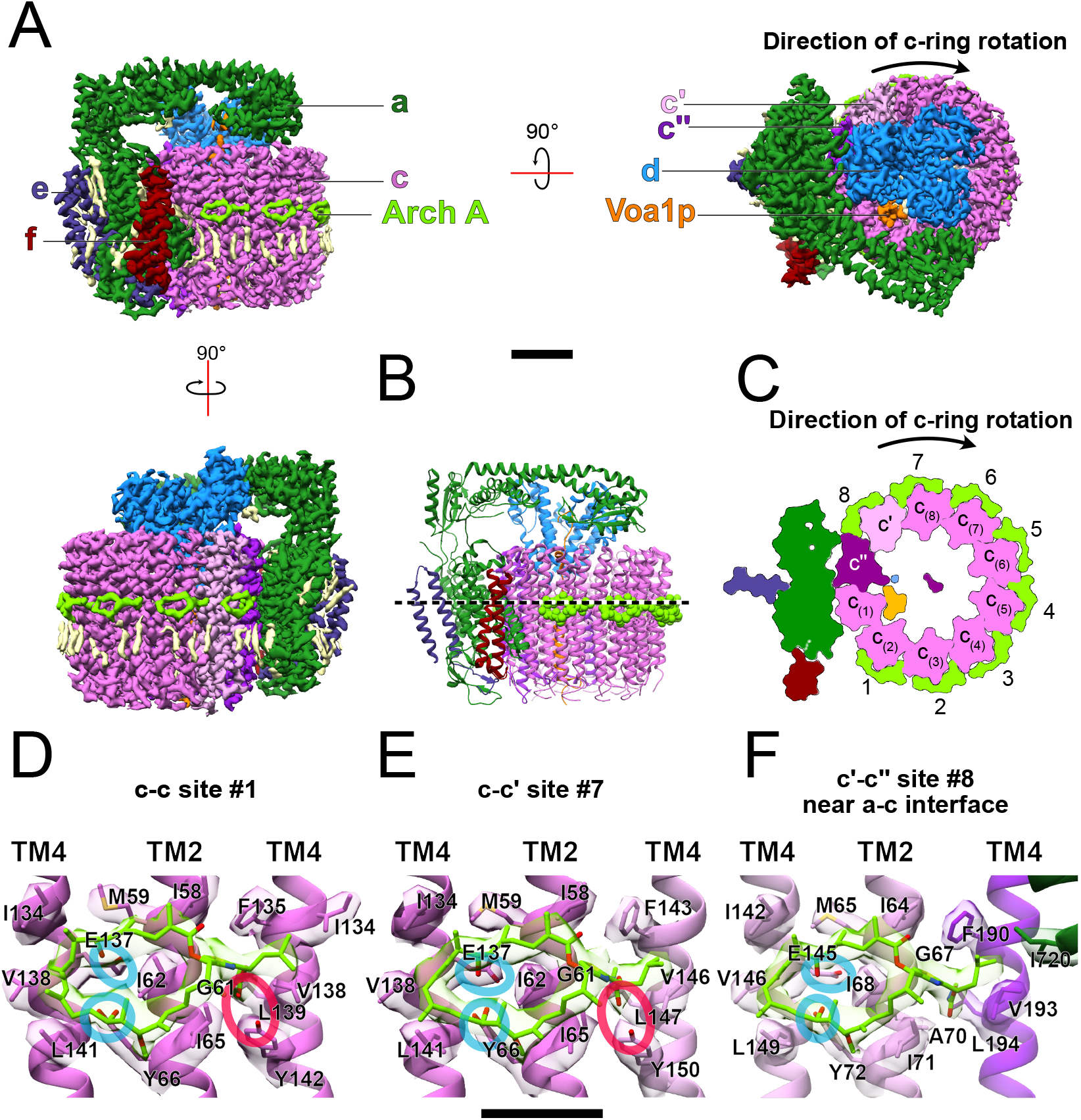
Structure of archazolid A bound to the yeast the V-ATPase V_O_ region. **A.** Cryo-EM map of archazolid A bound to the V _O_ region (three views). Scale bar, 25 Å. **B,** Atomic model for archazolid A bound to the yeast V _O_ region. **C,** A slice through the atomic model showing locations of eight archazolid A molecules. **D,** Atomic model showing a c-c site for archazolid A binding. **E,** Atomic model showing the c-c′ site for archazolid A binding. **F,** Atomic model showing thr c′-c′′site for archazolid A binding. Scale bar, 10 Å.

The archazolid density in the c′-c′′ site is well resolved, despite lacking one of the hydrogen bonds found in the c-c and c-c′ sites (Fig. S3H). Indeed, with the standard deviation of the map normalized to σ=1, archazolid molecules in the c-c sites become visible at σ=5 to 6, archazolid in the c-c′ site becomes visible at σ=6, and archazolid in the c′-c′′ site becomes visible at σ=6. This observation suggests that the hydrogen bond involving the amide oxygen in archazolid is not absolutely required for inhibitor binding. This hypothesis is supported by potent inhibition of V-ATPase by an archazolid analogue that lacks the thiazole group (Scheeff et al., 2020), and an increase in archazolid A potency when Tyr142 from subunit c was mutated (Bockelmann et al., 2010). Similarly, mutating Tyr66 from subunit c does not lead to archazolid A resistance (Bockelmann et al., 2010), suggesting that that hydrogen bond is not required for binding. Similar to bafilomycin, binding of archazolid around the c ring is expected to block rotation of the ring, thereby inhibiting V-ATPase activity.

### Comparison of bafilomycin and archazolid binding sites

Despite similarities in the structures of bafilomycin A1 and archazolid A (Fig. 1A and C, *left*) there is little overlap in the binding sites for these inhibitors. This lack of overlap is true at the c-c sites (Fig. 4A), the c-c′ site (Fig. 4B), and the c′-c′′ site (Fig. 4C). The small amount of overlap that does occur is near the Tyr142 of subunit c in the c-c and or Tyr 150 of subunit c′ in the c-c′ site that appear to participate in hydrogen bond formation with the 7′ hydroxyl group of bafilomycin and the 15′ hydroxyl group of archazolid. This overlap explains competitive binding seen between the two inhibitors (Bockelmann et al., 2010). In the c′-c′′ site (Fig. 4C), archazolid appears to bind with higher occupancy than bafilomycin. This difference is likely because bafilomycin binding is more dependent than archazolid binding on forming a hydrogen bond with the Tyr residue that is absent in the c′′ subunit. The different occupancy of bafilomycin and archazolid in the c′-c′′ is evident from viewing the maps at σ=4 with the standard deviation of the map normalized to σ=1 (Fig. 4C, *green and yellow densities*).

**Figure 4.**
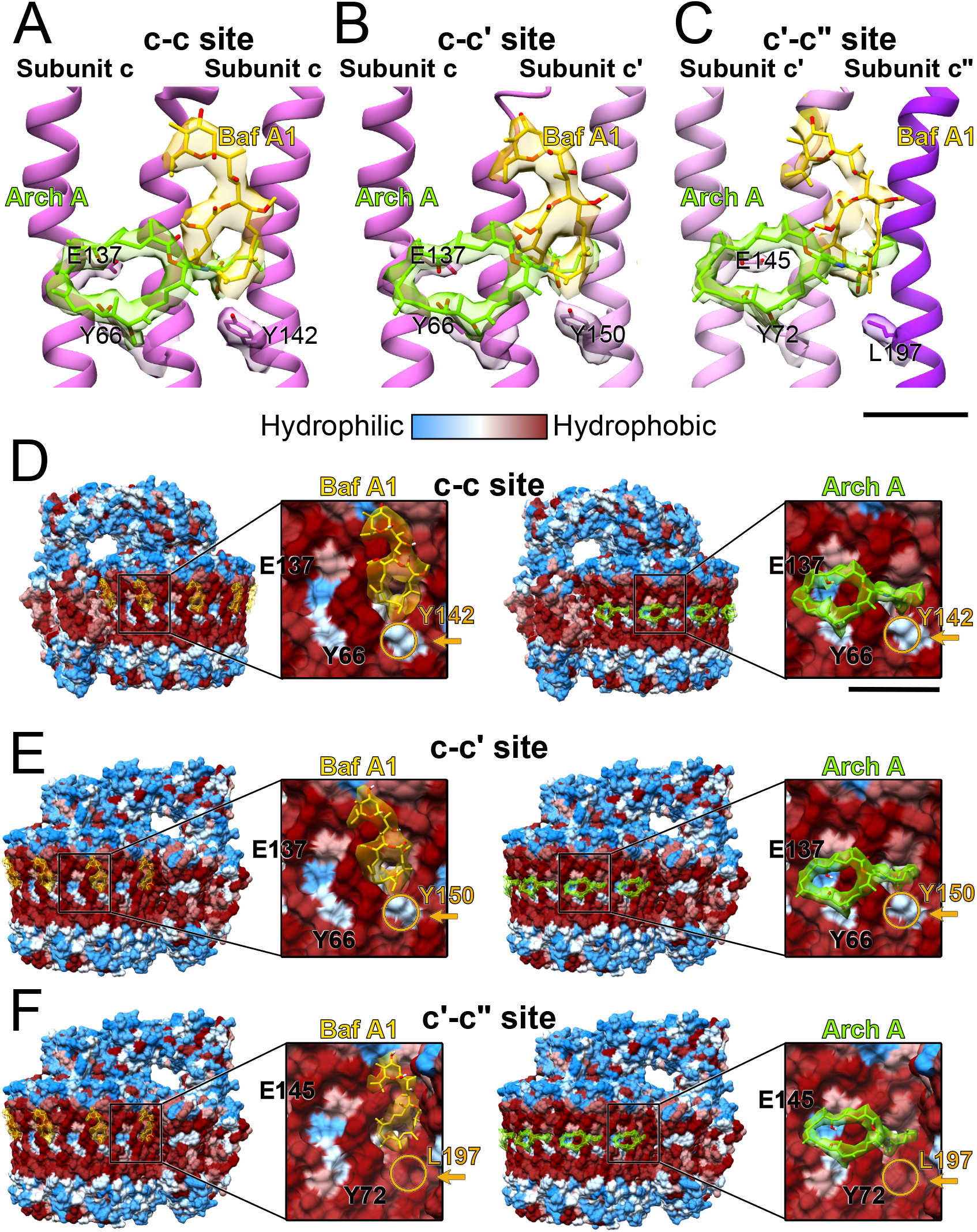
Comparison of the binding modes of bafilomycin A1 and archazolid. Atomic models showing the overlap of archazolid A (*green density and model*) and bafilomycin A1 (*yellow density and model*) in a c-c sites (**A**), the c-c′ site (**B**), and the c′-c′′ site (**C**). Scale bar, 25 Å. **D,** Kyte-Doolittle hydrophobicity with bafilomycin A1 (*yellow density, left*) or archazolid A (*green density, right)* shown in a c-c site (**D**), the c-c′ site (**E**), and the c′-c′′ site (**F**). Scale bar, 10 Å.

Other than the residues that likely participate in hydrogen bonds with the macrolides, binding of both inhibitors appears to be through van der Waals interactions due to close packing of the inhibitor onto the hydrophobic surface of the c ring (Fig. 4D to F). Putative hydrogen bonds with conserved Tyr and Glu residues (Fig. 4D to F, *blue patches*) appear to increase affinity but are not absolutely required for binding, as binding still occurs in the c′-c′′ site, which lack one of the Tyr residues involved in archazolid binding and the only Tyr residue involved in bafilomycin binding (Fig. 4D to F, *circled in orange*). This observation that the van der Waals interactions are sufficient for inhibitor binding is consistent with inhibitor binding still occurring when the hydrogen-bonding groups of the inhibitors are removed (Gagliardi et al., 1999, 1998; Scheeff et al., 2020) or when the residues involved in hydrogen bonding are mutated (Bockelmann et al., 2010).

The tyrosine residues involved in putative hydrogen bond formation are highly conserved and appear to be important for enzyme function (Bockelmann et al., 2010). Unlike bafilomycin A1, archazolid A binds this essential glutamate of the c and c′ subunits (residues Glu137, and Glu145, respectively). Direct binding to the proton carrying Glu residue is also seen in the ATP synthase inhibitors oligomycin (Symersky et al., 2012) and bedaquiline (Guo et al., 2021). With bedaquiline, markedly different affinities were observed between binding sites that involved just the c subunits and the binding sites that also involved the subunit a (Guo et al., 2021). However, only bafilomycin and not archazolid binding involves extensive contact with subunit a, and it is not clear whether this interaction contributes to its affinity. Remarkably, whereas the binding sites of the relatively small bedaquiline molecule on the ATP synthase c ring involve deep clefts in the structure and a rotation of the c ring to create these sites, the larger bafilomycin and archazolid binding sites do not involve clefts or rotation of the c ring (Fig. 4D to E). Instead, the flat surfaces from bafilomycin and archazolid form close contacts with flat surfaces on the V-ATPase c_8_c′c′′ ring surface. Not requiring a cavity in the c ring for binding means that there are more possible binding sites for inhibitors, allowing for binding sites with only minimal overlap for bafilomycin and archazolid. Binding to the flat surface also indicates that other macrolides could bind different parts of the c_8_c′c′′ ring surface, suggesting a large chemical space available to V-ATPase inhibitors. The diversity of available binding sites for macrolide inhibitors of V-ATPase suggests that numerous different inhibitors may be identified, each with different pharmacokinetic or physicochemical properties.

## Methods

### Protein purification

Intact V-ATPase (Benlekbir et al., 2012) and the V_O_ complex (Mazhab-Jafari et al., 2016) were purified as previously described with some modifications. Intact V-ATPase was purified from the *Saccharomyces cerevisiae* strain SABY31 (Vma1p-3×FLAG,ΔStv1p) while the V_O_ complex was purified from *S. cerevisiae* strain CACY1 (Vph1p 3×FLAG). Yeast were grown at 30 °C on YPD agar plates (1 % [w/v] yeast extract, 2 % [w/v] peptone, 2 % [w/v] D-glucose, 1.5 % [w/v] agar) and used to inoculate two 100 mL cultures of liquid YPD medium. These cultures were grown at 30 °C overnight (>12 h) with shaking at 220 rpm. The cultures were then used to inoculate YPD (11 L) in a Microferm fermenter (New Brunswick Scientific), which was incubated for two days (>36 h) with stirring at 300 rpm and aeration at 34 cubic feet per h.

All cell harvest and protein purification steps were performed at 4 °C. Cells were harvested by centrifugation at 4,000 ×g for 15 min. Cell pellets were resuspended in lysis buffer at 1 mL/g cell pellet (10 mM Na_2_HPO_4_ and 2 mM KH_2_PO_4_ pH 7.4, 140 mM NaCl, 3 mM KCl, 8 % [w/v] sucrose, 2 % [w/v] D-sorbitol, 2 % [w/v] D-glucose, 5 mM 6-aminocaproic acid, 5 mM benzamidine, 5 mM ethylenediaminetetraacetic acid [EDTA], 0.001 % [w/v] phenylmethanesulfonyl fluoride [PMSF]). Yeast cells were lysed in a bead beater with 0.5 mm glass beads (BioSpec), which was cooled with ice, using four cycles of 1 min of bead-beating followed by 1 min of cooling. Cell debris was removed by centrifugation at 3,500 ×g for 10 min. Cell membranes were then collected by centrifugation at 110,000 ×g for 40 min, resuspended in lysis buffer at 0.5 mL/g original cell pellet, and divided into 30 to 45 mL fractions for flash freezing in liquid nitrogen and storage at −80 °C.

Membranes were thawed and solubilized with 1 % (w/v) n-dodecyl β-D-maltoside (DDM) and insoluble material was removed by centrifugation at 150,000 ×g for 70 min. Solubilized membranes were filtered with a 0.45 μm filter and bound to 0.8 mL of anti-FLAG M2 affinity gel (Sigma) previously equilibrated in DTBS (50 mM Tris-HCl pH 7.4, 150 mM NaCl, 0.05 % [w/v] DDM) in a disposable column. The beads were washed with ten bed volumes of DTBS and V-ATPase was eluted with 1.5 mL of DTBS buffer containing 150 μg/mL 3×FLAG peptide followed by 0.5 mL of DTBS without peptide. Purified V-ATPase in buffer containing DDM was used in enzyme assays.

To partially exchange the V_O_ complex into a buffer containing the detergent glyco-diosgenin (GDN) for cryo-EM, protein was concentrated to ~100 μL with a VivaSpin6 100 kDa cutoff concentrator centrifuged at 1,000 ×g. The sample was diluted with 4 mL of GTBS buffer (50 mM Tris-HCl pH 7.4, 150 mM NaCl, 0.004 % [w/v] GDN) followed by concentration in the same concentrator to ~1 mL. For cryo-EM of the bafilomycin-bound structure, a 5 mM stock of bafilomycin A1 (Cayman Chemistry) in DMSO (Sigma) was added to the sample to a final concentration of 50 μM (1% [v/v] DMSO). Similarly, for cryo-EM with archazolid A (isolated from *Cystobacter* sp. strain Cbm 20), a 10 mM stock of inhibitor was added to the sample to a final concentration of 100 μM (1% [v/v] DMSO). For both inhibitors, the enzyme-inhibitor mixture was incubated on ice for 45 min. Samples were then further concentrated with the VivaSpin6 concentrator for 5 min at 1,000 ×g, and then concentrated to a final protein concentration of ~3 mg/mL (25 μL volume) with a VivaSpin500 100 kDa cutoff concentrator for 15 min at 12,000 ×g.

### Enzyme assays

Enzyme-coupled ATPase activity assays with V-ATPase were performed as described previously (Bueler and Rubinstein, 2015; Lötscher et al., 1984; Vasanthakumar et al., 2019) with some modifications. Assays were performed in a 96-well plate with a total reaction volume of 160 μL of assay buffer (50 mM Tris-HCl pH 7.4, 3 mM MgCl_2_, 0.2 mM NADH disodium salt, 3.2 units pyruvate kinase, 8 units L-lactic dehydrogenase, and 0.02% [w/v] DDM, 100 μg/mL soybean asolectin [Sigma]). V-ATPase was measured before experiments and the enzyme was used at final concentrations of 0.79 to 6.4 nM. Inhibitors in DMSO were added from stock solutions, with the final concentration of DMSO in the reaction maintained at 1% (v/v). Plates were incubated at 37 °C and reactions were initiated by addition of ATP disodium salt and phosphoenol pyruvic acid monopotassium salt to 2 mM and 1 mM, respectively. Absorbance at 340 nm was monitored at 37 °C for 10 min, and the linear portions of the absorbance versus time curve was used to calculate enzyme activity. For specific activity measurements, absorbance at 340 nm was converted into NADH concentration with a standard curve. Individual titrations of V-ATPase activity with inhibitors were fit to the Hill equation using non-linear regression with the software *R* (R Core Team, 2021). The arithmetic mean of the IC_50_ values from each titration and standard deviation are reported. Dose-response plots were generated with *ggplot2* (Wickham, 2016).

### Cryo-EM and image analysis

Cryo-EM specimens were prepared using home-made nanofabricated holey gold grids (Marr et al., 2014) with ~1.5 μm holes. Grids were glow-discharged in air for 120 s and 1.5 μL of sample was applied to the grids, blotted for 2 s with a Leica EM GP2 freezing device at 80 to 90 % relative humidity, and frozen in liquid ethane. Sample screening for grid optimization was done with a FEI Tecnai F20 electron microscope operating at 200 kV and equipped with a Gatan K2 Summit camera. Final data collection was performed with a Titan Krios G3 electron microscope (Thermo Fisher Scientific) operating at 300 kV and equipped with a Falcon 4 camera (bafilomycin dataset) or a prototype Falcon 4i camera (archazolid dataset). Data collection was automated with the *EPU* software package. Movies consisting of 29 exposure fractions were collected at a nominal magnification of 75,000 ×, corresponding to a calibrated pixel size of 1.03 Å. For the bafilomycin A1 and archazolid A datasets, total exposures of ~39 electrons/Å^2^ and ~45 electrons/Å^2^ were used, respectively, with exposure rates of 5.42 and 6.37 electrons/pixel/s, respectively.

Except where noted, image analysis was performed with *cryoSPARC* v.3 (Punjani et al., 2017). For the bafilomycin dataset, 3,960 movies were aligned with *MotionCor2* (Zheng et al., 2017) using a 7×7 grid. For the archazolid dataset, 4,094 movies were aligned in patches in *cryoSPARC*. For both datasets, contrast transfer function (CTF) parameters were estimated in patches. Templates for particle selection were generated by 2D classification of particle images following automatic selection of particles with the blob picker algorithm and datasets of particle images were extracted in 300×300 pixel boxes. This process provided 1,278,735 particle images for the bafilomycin dataset and 1,723,337 particle images for the archazolid dataset. The images were then subjected to 2D classification and 707,137 particle images from the bafilomycin dataset and 1,183,038 particle images from the archazolid dataset were selected for further analysis.

For the bafilomycin dataset, image parameters were converted to *Relion* 3.0 .star file format with the *pyem* package (https://zenodo.org/record/3576630#.X5_WHFX7TIU), particle images were downsampled to 256×256 pixel boxes, and motion was corrected for individual particles in *Relion* with the Bayesian polishing algorithm (Zivanov et al., 2019). Images were imported back into *cryoSPARC* v.3, and several rounds of *ab initio* 3D classification followed by heterogeneous refinement were performed. The remaining images were then subjected to non-uniform refinement, global CTF refinement, and local CTF refinement, followed by an additional round of non-uniform refinement with individual particle CTF refinement. This process produced a map at 3.2 Å resolution.

Local motion correction (Rubinstein and Brubaker, 2015) was performed for particle images from the archazolid dataset. These images were then subjected to several rounds of *ab initio* 3D classification and heterogeneous refinement. A 3D class corresponding to the archazolid-bound V_O_ complex was refined with non-uniform refinement (Punjani et al., 2020), global CTF refinement, and local CTF refinement, followed by an additional round of non-uniform refinement and individual particle CTF refinement. This process produced a map at 2.8 Å resolution.

## Acknowledgements

KAK was supported by a Canada Graduate Scholarship from the Canadian Institutes of Health Research (CIHR), an Ontario Graduate Scholarship, and a McCuaig-Throop Bursary from the University of Toronto. JLR was supported by the Canada Research Chairs program. This research was supported by Canadian Institutes of Health Research grant PJT166152 (JLR). Cryo-EM data were collected at the Toronto High-Resolution High-Throughput cryo-EM facility, supported by the Canada Foundation for Innovation and Ontario Research Fund.

---

**Supplementary Figure 1.**
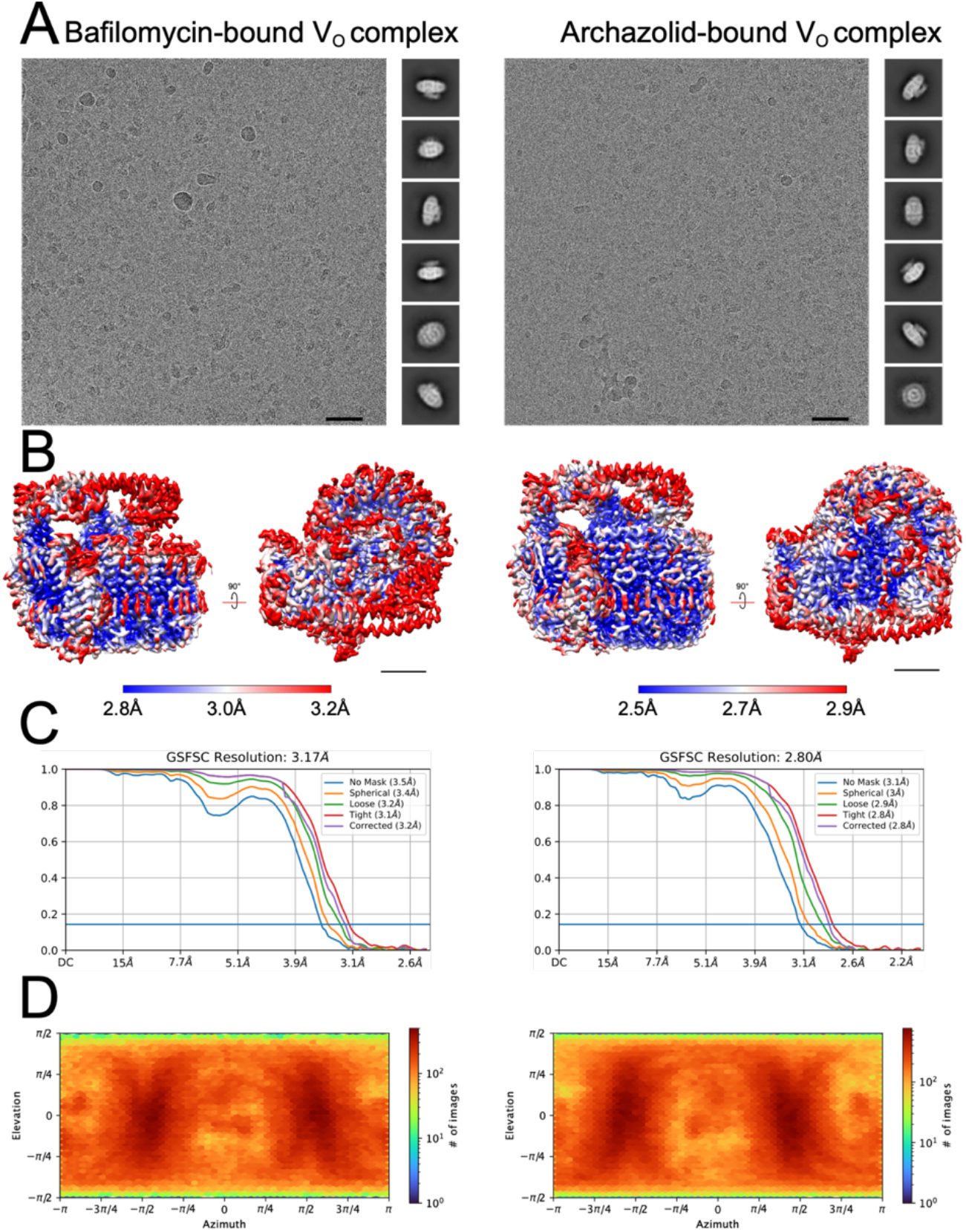
Cryo-EM data and validation. **A,** Representative micrographs and 2D class average images of the bafilomycin A1 (*left*) and archazolid A (*right*). Scale bar, 100Å. **B,** Local resolution maps for the bafilomycin A1-bound V_O_ structure (*left*) and the archazolid A-bound V_O_ structure (*right*). Scale bar, 25Å. **C,** Fourier Shell correlations of the bafilomycin A1-bound V_O_ structure (*left*) and the archazolid A-bound V_O_ structure (*right*). **D,** Orientation distribution of particle images for the bafilomycin A1-bound V_O_ structure (*left*) and the archazolid A-bound V_O_ structure (*right*).

**Supplementary Figure 2.**
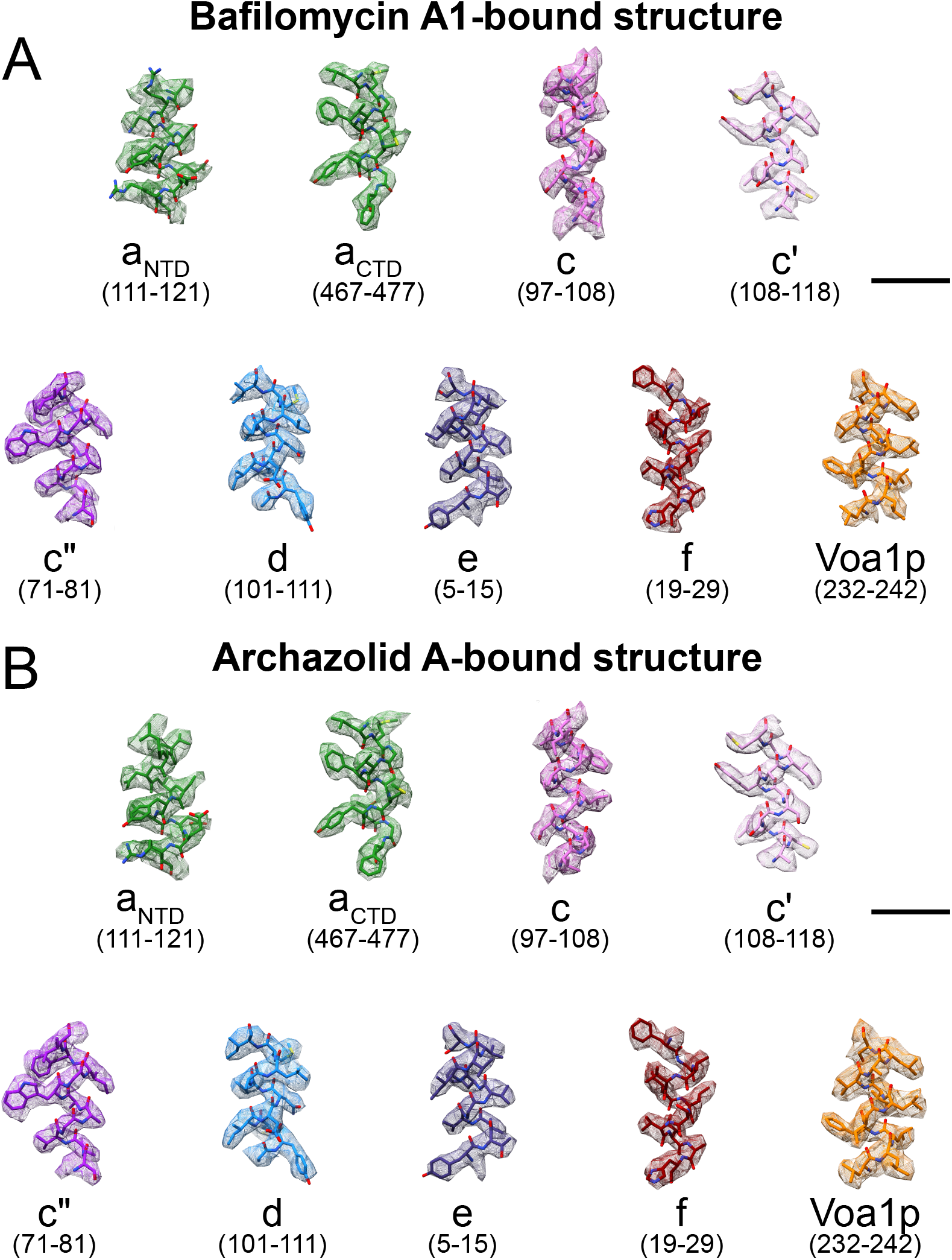
Model-in-map fit examples. Examples of atomic model fit in the experimental cryo-EM density map for the bafilomycin A1-bound V_O_ structure (**A**) and archazolid A-bound V_O_ structure (**B**).

**Supplementary Figure 3.**
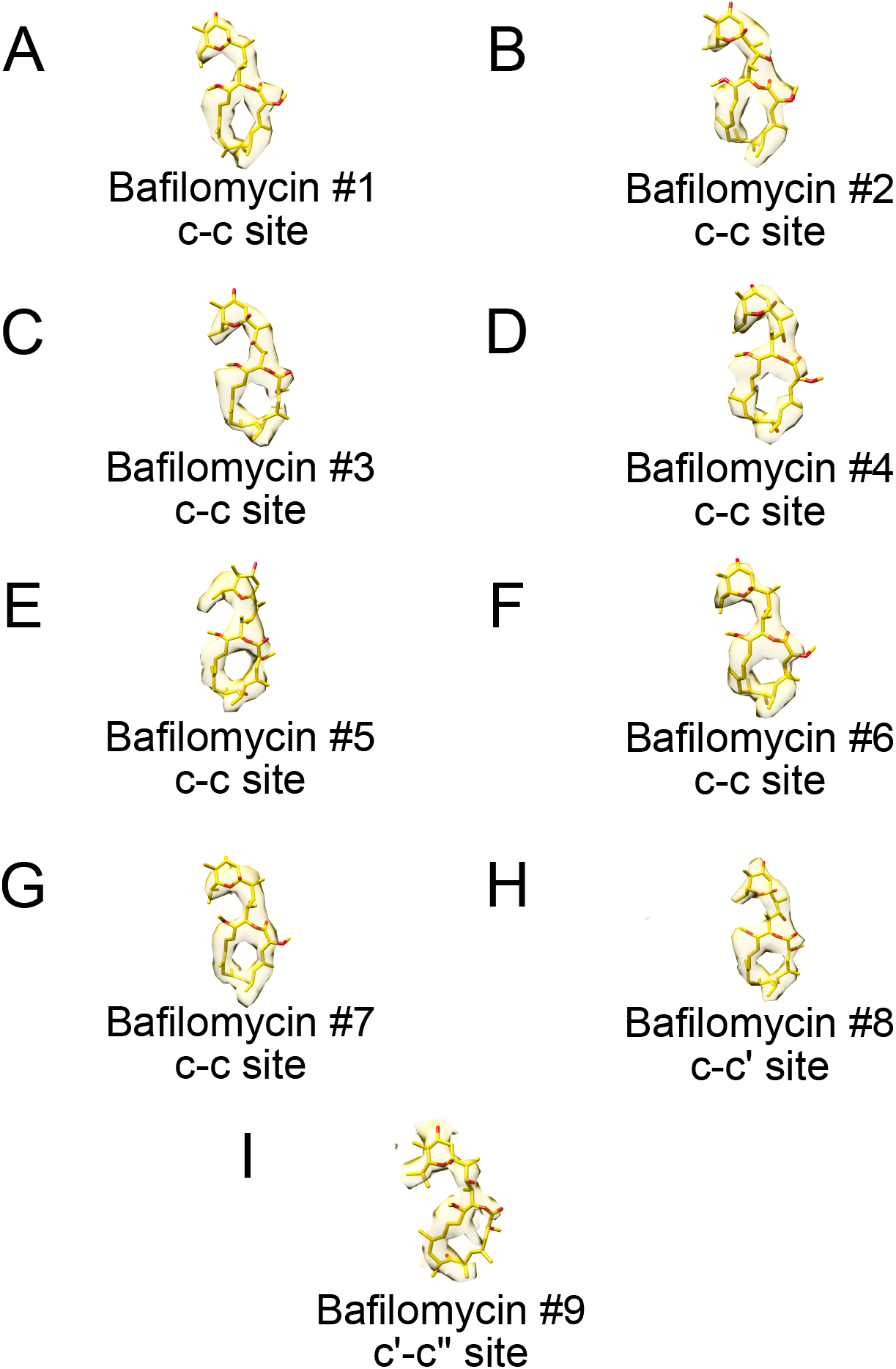
Individual bafilomycin A1 densities and atomic models. **A,** Bafilomycin A1 #1. **B,** Bafilomycin A1 #2. **C,** Bafilomycin A1 #3. **D,** Bafilomycin A1 #4. **E,** Bafilomycin A1 #5. **F,** Bafilomycin A1 #6. **G,**Bafilomycin A1 #7. **H,** Bafilomycin A1 #8. **I,** Bafilomycin A1 #9. Bafilomycin numbering corresponds to Fig. 2.

**Supplementary Figure 4.**
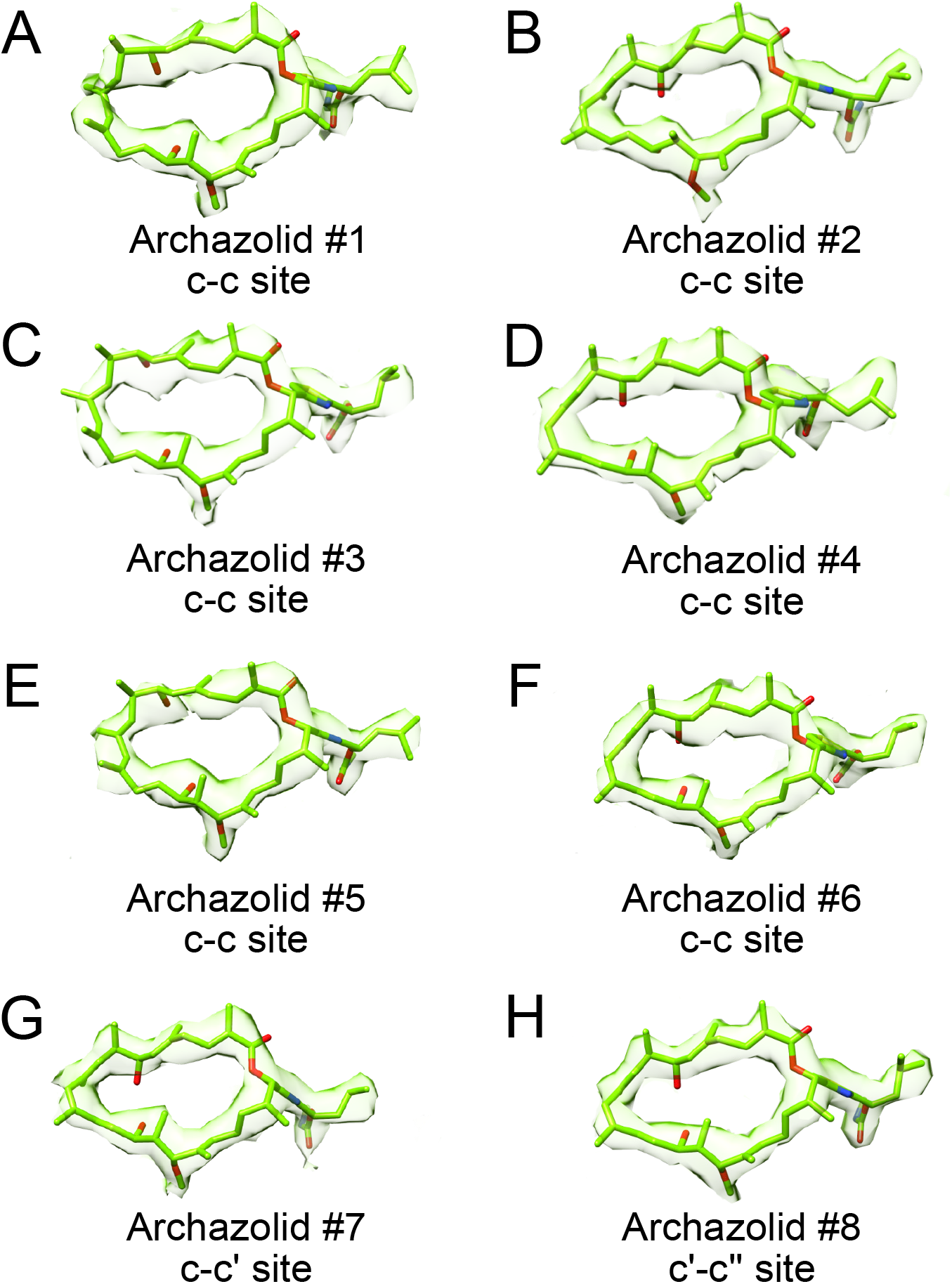
Individual archazolid A densities and atomic models. **A,** Archazolid A #1. **B,** Archazolid A #2. **C,** Archazolid A #3. **D,** Archazolid A #4. **E,** Archazolid A #5. **F.** Archazolid A #6. **G,** Archazolid A #7. **H,** Archazolid A #8. Archazolid numbering corresponds to Fig. 3.

